# Concurrent Supervised-Unsupervised Representative Subspace Clustering (CSU-RSC)

**DOI:** 10.64898/2026.09.09.750386

**Authors:** Youngjo Song, Shiva Mirzaeian, Pablo Andrés-Camazón, Vince Calhoun, Jiayu Chen, Armin Iraji

## Abstract

The biological heterogeneity of psychotic disorders (PDs) has motivated the identification of psychosis imaging neurosubtypes (PINs), which aims to stratify the PD population into subgroups with similar neurobiological underpinnings. Yet the influence of sex, although well documented in the literature, has not been well counted in the existing subtyping paradigm. Here, we introduce *concurrent supervised-unsupervised representative subspace clustering* (CSU-RSC), a general functional neurosubtyping framework that jointly estimates cluster-specific subspaces and a partition while softly incorporating a supervised label, aiming to identify sex-dominant subgroups based on differences in the spatial organization of brain networks. This soft supervising design differentiates CSU-RSC from unsupervised approaches that completely ignore a label as well as from supervised approaches that exclusively analyze each label group separately. On simulated data, CSU-RSC recovered the ground truth of subgroups more accurately than its unsupervised counterpart. Applied to resting-state fMRI from 1,239 individuals with psychosis, CSU-RSC identified two sex-dominant PINs that replicated across discovery and validation sets, and showed significant differences in clinical characteristics, including cognitive impairment and symptom severity, with the most cognitively impaired subtype being female-dominant. Projecting controls onto the patient-derived subspaces showed that the female-dominant pattern did not persist in controls, suggesting psychosis-specific sex heterogeneity. Overall, by incorporating sex as an explicit source of information, CSU-RSC highlights the value of sex for resolving biological heterogeneity and provides a basis for more reproducible and biologically informative patient stratification.

## Introduction

Psychotic disorders (PD) often converge on a shared set of symptoms, yet are biologically heterogeneous, influenced by many different genetic and environmental factors [1–3]. For instance, in the psychosis spectrum, schizophrenia (SZ), schizoaffective disorder (SAD), and bipolar disorder with psychosis (BP) share overlapping symptoms (e.g., delusions, hallucinations, disorganized thinking), blurring their diagnostic boundaries [4–6]. Yet within and across these diagnoses categories, notable differences in neurobiology have been observed, presenting divergent brain structural and functional alterations [7, 8] and variable treatment response [9]. These findings clearly point to significant heterogeneity in the complex biological underpinnings of PD. To address this, identification of psychosis imaging neurosubtypes (PINs)—stratifying the PD population based on their neurobiological profiles, such as fMRI-based measures, rather than just clinical symptoms—has recently emerged as a way to define more homogeneous, treatment-relevant subgroups [10–12].

Intrinsic connectivity networks (ICNs) estimated from resting-state fMRI (rsfMRI) data provide a natural basis for such subtyping. ICNs, typically estimated by independent component analysis (ICA), are functionally coherent sources whose engagement reflects ongoing cognitive and mental states [13, 14]. Their spatial topography varies systematically across individuals (cf. [15, 16]; see also [17–19]), reflecting individual differences in functional network organization, and such variation is clinically relevant in PDs [20, 21], with links to symptom presentation and illness course (e.g., [22, 23]). A central challenge is to characterize such individual variation in a way that preserves the structured heterogeneity that may define PINs. Most existing subtyping methods reduce each subgroup to a single summary: centroid-based approaches (e.g., k-means, hierarchical clustering) represent a subgroup by one representative profile of brain activities (e.g., [24–26]), while discriminative approaches (e.g., HYDRA [27], CHIMERA [28]) represent it by the direction in which patients depart from a reference group. In both, the subgroup collapses to a single point or direction in the high-dimensional feature space, so that within-subgroup variability (whether noise or meaningful individual differences) is treated only as a scalar deviation from that centroid/reference rather than as structure.

In contrast, subspace-based clusterwise decomposition (e.g., clusterwise independent component analysis (C-ICA) [29], clusterwise simultaneous component analysis (C-SCA) [30], clusterwise Parafac [31]) can define each subgroup by a shared low-dimensional subspace of brain activities (i.e., the space spanned by a set of ICNs) rather than a single centroid. This is particularly relevant for rsfMRI, where biologically meaningful heterogeneity can manifest as differences in the spatial organization of multiple ICNs. Under a subspace model, between-subtype heterogeneity is represented by distinct network bases (thus capturing heterogeneity from multiple ICNs), whereas within-subtype heterogeneity is accommodated through individual-specific expressions (i.e., mixing) of a shared set of ICNs. Consequently, by separating two sources of variation, subtypes are distinguished by genuine differences in network organization rather than by individual variability within a common network structure.

One notable aspect is that sex, as a biological factor, contributes significantly to heterogeneity in psychosis. Sex differences in psychosis are well documented across symptom profiles [32], illness course [33], and brain function [34]. Despite this, sex has not been explicitly incorporated into existing subspace-based decomposition frameworks. More broadly, in typical subtyping, network structures are typically estimated without leveraging known demographic or biological factors that may organize heterogeneity, potentially obscuring structured differences along these axes. Incorporating such information during subtyping may therefore enable a more targeted characterization of biologically meaningful heterogeneity.

To this end, we introduce *concurrent supervised-unsupervised representative subspace clustering* (CSU-RSC), a general framework that softly incorporates an arbitrary supervised label into an arbitrary clusterwise decomposition (e.g., ICA). Its defining feature is that tunable parameters flexibly span an entire continuum: at one extreme the label is ignored, recovering purely unsupervised clustering (i.e., C-ICA); at the other, each label group is fully separated, recovering label-wise clustering (i.e., stratification); and in between, the label is only partially enforced. Using simulations with known ground truth and empirical psychosis rsfMRI data with sex as the supervised label, we demonstrate the framework’s validity and its ability to derive reproducible, sex-dominant PINs.

## Methods

### Concurrent Supervised-Unsupervised Representative Subspace Clustering

Concurrent supervised-unsupervised representative subspace clustering (CSU-RSC) algorithm softly incorporates a supervised label into a subspace-based clusterwise decomposition procedure. CSU-RSC generalizes the clusterwise decomposition methods in two ways: (i) the subspace estimator is treated as an interchangeable module, so approach such as an linear latent-variable models (e.g., PCA, ICA) or deep representation-learning models (e.g., autoencoder) can be used, which enhances scalability; and (ii) a supervised label is used to constrain, but not fully dictate, the partitioning.

#### Notations

Let *X_i_ (i* = *1,* …, *I*) denote the *V* x *T* data matrix of subject *i* (V: number of voxels, *T*: time points). Each subject carries a known supervised label *u* G {1, …, *U}* where U denotes the number of label categories (e.g., *U = 2* for sex, male or female). Subjects are partitioned into *R*= *U×W* mutually exclusive partitions, where *W* is the number of unsupervised sub-partitions estimated per labeled group. Note that CSU-RSC does not restrict each partition to contain only subjects with its corresponding supervised label. Rather, each partition is (softly) encouraged to be dominated by, but not exclusive to, that label. Each partition *r* = *(u,* w) is represented by a shared subspace basis *S^(r)^*(*V×Q*,*Q* is number of bases/components) obtained by decomposing the concatenated data of its members. Let *P* (Z x *R*) be the binary partition matrix, with *P_ir_* = 1 i.f.f. subject *i* is assigned to partition *r;* otherwise *P_ir_* = 0. Writing 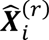 for the reconstruction of *X_i_* from the subspace of partition *r*, CSU-RSC optimizes the following reconstruction loss:

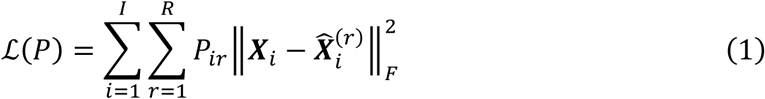

In the linear case, 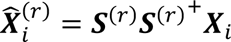 where *S^(r)+^* is the pseudo-inverse of *S(r)*; more generally, 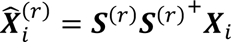 may be produced by any encoder-decoder pair (e.g., an autoencoder) per partition.

#### Algorithm

CSU-RSC generally minimizes reconstruction loss *ℒ* through the constrained optimization detailed below (see Figure 1 for an overview). However, it is noteworthy that CSU-RSC does not minimize reconstruction error alone: it softly incorporates the label (here, sex) into the optimization, guiding the partitioning toward label-relevant structure while still allowing subjects of either label to group together. Below is a detailed procedure:

1. **Initialization.** Each subject is assigned to a random sub-partition *within its own labeled group*. Seeding partitions with same-label subjects guiding the initial subspaces toward label-specific structure.
2. **Subspace estimation.** For each partition *r,* subject fMRI time-series data are decomposed to obtain group-level subspace bases, **S^(r)^**. The decomposition can use any method that yields a low-dimensional latent representation (e.g., PCA, ICA, or autoencoder).
3. **Reassignment.** Each subject is reassigned to the partition minimizing its reconstruction error (i.e., projection residual) 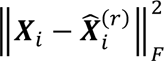. Reassignment is unrestricted: a subject may be assigned to any partition regardless of supervised label; label mismatch is handled subsequently by the subsequent refinement mechanism.
4. **Refinement.** Before the next subspace estimation, two mechanisms temporarily remove subjects to refine partition-specific structure: (1) pruning by affinity and (2) dropout by supervised label. *(Pruning by affinity)* For pruning, we compute a partition affinity score, *PASi* = (*bi … a_i_)/max^(^a_i_,bt),* where *a_i_* is the residual to the assigned partition and *bi* the smallest residual to any other partition (analogous to the silhouette value [35]). Subjects with PASj below the cohort mean (i.e., those weakly or ambiguously assigned) are excluded for this iteration. *(Dropout by supervised label)* Among retained subjects whose supervised label differs from that of their assigned partition, each is dropped at random with a predefined probability. Pruning and dropout are per-iteration only; removed subjects re-enter the pool for reassignment in the next iteration. Both refinement steps and their rates are configurable and can be disabled depending on researcher’s choices.
5. **Convergence and selection.** Steps 2–4 repeat until convergence, defined as no improvement in the *ℒ* exceeding a tolerance of 10⁻⁶ for a fixed number of consecutive iterations (e.g., 10).
6. **Multi-run consensus selection.** For algorithmic stability, CSU-RSC is run M times with random initializations, and a single representative run is selected. For every pair of runs, we compute the adjusted Rand index (ARI) [36] and retain the run with the highest mean ARI with all other runs as the final solution: i.e., the solution most representative of the dominant, reproducible partitioning across random initializations.

**Figure 1.**
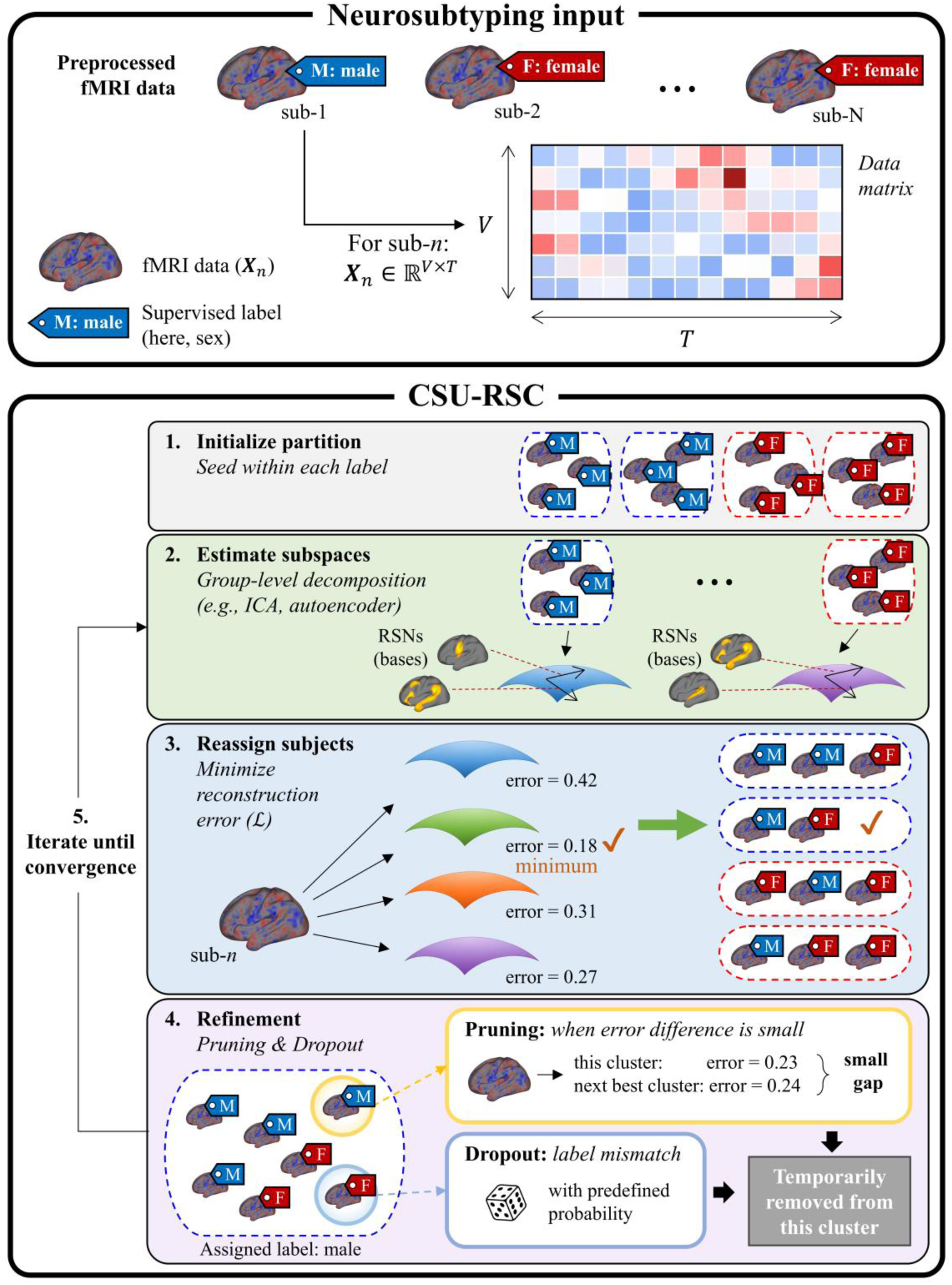
Overview of CSU-RSC. **(Top)** Neurosubtyping input. Each subject’s preprocessed fMRI data *X_n_* (V voxels x T time points, *n*-th subject) is paired with a supervised label (here, sex; M = male, blue; F = female, red). **(Bottom)** The optimization loop of the CSU-RSC procedure. (1) Subjects are initialized into partitions (clusters) seeded within each label, so that each partition initially contains subjects of a single label. (2) Each partition’s subspace is estimated by group-level decomposition (e.g., ICA) of the fMRI data of its member subjects, spanned by a set of resulting bases (e.g., ICs). (3) Each subject is reassigned to the partition whose subspace minimizes its reconstruction error, regardless of its supervised label. (4) Refinement temporarily removes subjects through two mechanisms: pruning, which removes weakly assigned subjects (those with a small error gap to the next-best cluster) regardless of supervised label, and dropout, which removes label-mismatched subjects with a predefined probability. Removed subjects re-enter at the next iteration, so partitions remain label-dominant but not label-exclusive. In the final assignment, refinement is omitted so that all subjects are retained. Steps 2–4 iterate until convergence, defined as no improvement in total reconstruction error for a fixed number of consecutive iterations (patience-based stopping). Note that the label-separated partitions depicted in step 2 are schematic and reflect only the initial iteration; after reassignment in subsequent iterations, partitions may contain subjects from both labels. See Methods for detailed procedures.

#### Implementation

For the analysis in this paper, we made the following specific choices within the CSU-RSC framework. We used sex as the supervised label, allowing the partitioning to be guided by sex while estimating two unsupervised sub-partitioning within each sex (*W* = 2), i.e., four subtypes in total. Accordingly, the algorithm encourages the four subtypes to organize into two male-dominant and two female-dominant groups, although this structure is guided rather than enforced. For decomposition, we used group PCA rather than group ICA to estimate each partition’s subspace. Because group ICA procedure rotates the PCA-derived basis without changing the subspace it spans, the two decompositions yield the same partition subspace, and therefore the same reconstruction error and cluster assignments. Thus, group PCA was chosen for computational efficiency, avoiding the additional basis rotation step at each iteration. Independent components (ICs) are estimated post hoc using infomax ICA once the final partitions are fixed, for interpretability. We fixed the number of components per partition to Q = 10, so that a compact set of components captures the reliable structure while reducing overfitting [37]. For refinement, we applied pruning based on the PAS, removing weakly or ambiguously assigned subjects at each iteration, and set dropout to 100%, removing all subjects whose supervised label differed from that of their assigned partition. Note that even at 100% dropout the partitions are not constrained to be single-sex, since dropped subjects re-enter the pool at the next iteration and the supervised constraint is fully relaxed at the final assignment. We performed 30 runs (i.e., M = 30) and selected the most reproducible one (i.e., the run with the highest mean ARI).

### Alternative clustering algorithms

To compare our proposed CSU-RSC algorithms with alternative approaches, we implemented other clustering methods commonly used for subtyping.

#### C-ICA algorithm

Clusterwise ICA (C-ICA) [29] can be viewed as the fully unsupervised extreme of CSU-RSC, in which the supervised label plays no role: no label-based constraint is imposed at initialization or reassignment, and no dropout is applied. C-ICA partitions subjects such that each cluster is characterized by its own brain-activity subspace, spanned by a set of shared independent sources (i.e., ICs). Similar to CSU-RSC, since group PCA and group ICA span the same subspace, we replaced the group ICA step with group PCA for computational efficiency. C-ICA minimizes the total reconstruction error through an alternating least squares procedure that iterates two steps until convergence: (1) given the current partition, the data of each cluster are temporally concatenated and decomposed by group PCA to obtain that cluster’s subspace; and (2) given the subspaces, each subject is reassigned to the cluster whose subspace best reconstructs its data. Because the procedure may converge to a local optimum, it is run from multiple random initializations (here, 30) and the solution with the lowest reconstruction error is retained.

#### HYDRA clustering algorithm

We also compared CSU-RSC to HYDRA [27], a semi-supervised method that discovers disease subtypes while separating patients from a reference group. Unlike the fully unsupervised C-ICA method, HYDRA requires both patients and cognitively normal controls CN as input: it groups patients by the direction in which they deviate from the CN. Specifically, it fits multiple linear max-margin hyperplanes that jointly form a convex polytope separating patients from CN, and each patient is assigned to the face that best discriminates it from the controls, so that each face captures a distinct pathological pattern. The classifier and the subject assignments are estimated by iterative alternating optimization. Since the objective is non-convex, the procedure was repeated over multiple initializations and cross-validation folds, and the final subtypes were obtained by consensus clustering of the resulting solutions (see [27] for details). In this work, we used the publicly available implementation (https://github.com/evarol/HYDRA), with default parameter settings under 5-fold cross-validation.

To keep the comparison within the domain of intrinsic functional networks, subject-specific multi-scale functional network connectivity (msFNC) served as the input features for HYDRA. Subject-specific time courses for the 105 multi-scale intrinsic connectivity networks of the NeuroMark 2.2 template [38, 39] were estimated from each subject’s fMRI data using spatially constrained ICA (MOO-ICAR, multi-objective optimization ICA with reference [40, 41]). Prior to connectivity estimation, the network time courses were postprocessed by detrending (linear, quadratic, and cubic), regressing out the six motion parameters and their derivatives, despiking via a third-order spline fit (median-absolute-deviation threshold = 2.5), and bandpass filtering (0.01–0.15 Hz). msFNC was then computed as the Pearson correlation between every pair of network time courses, yielding a 105 × 105 connectivity matrix per subject. The upper-triangular entries were vectorized and reduced to 10 PCs, matching the number of components (Q = 10) used in the C-ICA and CSU-RSC analyses.

#### CHIMERA clustering algorithm

We also implemented CHIMERA [28], a probabilistic clustering method that, like HYDRA, discovers patient subtypes relative to a CN reference group but does so through distribution matching rather than max-margin separation. CHIMERA treats CN and patients as two sets of points in an imaging feature space and models the disease as a set of regularized linear transformations mapping the control set onto the patient set, each presumably representing a distinct pathological direction. The transformations are estimated by a variant of coherent point drift within an expectation-maximization framework, and each patient is assigned to the transformation most likely to have generated it. Because the objective is non-convex, the procedure was repeated 30 times from random initializations, and the most reproducible solution (i.e., the run with the highest mean ARI with all other runs) was retained as the final subtyping. We used the publicly available implementation (https://github.com/anbai106/CHIMERA) with the same msFNC input features as described for HYDRA above (reduced to 10 PCs).

A notable feature of CHIMERA is that covariates are incorporated through a multi-kernel matching scheme, softly mitigating their confounding influence during distribution matching rather than regressing them out beforehand. We included age and site as covariate, but sex was not included so that any sex-related organization of the disease effect could be expressed in the transformations.

#### K-mean clustering

Finally, we implemented a K-means clustering procedure. Using the same input features as HYDRA and CHIMERA (msFNC reduced to 10 PCs), we clustered patients into four clusters (via built-in K-means function in MATLAB). The clustering repeated 30 times, and the run with the lowest loss was selected.

### Discovery–validation design and out-of-sample subtype assignment

To evaluate the replicability of the identified psychosis subtypes, the 1,239 patients were randomly divided into discovery and validation sets using an 80:20 split, yielding 992 discovery and 247 validation subjects, respectively. Subtyping was performed on the discovery set using CSU-RSC, C-ICA, HYDRA, CHIMERA, or K-means, and validation subjects were subsequently assigned according to the subtype representations derived from the discovery set.

Because the methods use different features and procedures to identify PINs, the assignment procedure varied by method. For the subspace-based methods, CSU-RSC and C-ICA, each discovery subtype was represented by its ICN subspace; each validation subject’s fMRI data was projected onto all four discovery-derived subspaces and assigned to the subtype with the minimum reconstruction error (see eq. 1). For HYDRA, CHIMERA, and K-means, the exemplar for each PIN was calculated for each discovery cluster as the mean of the input features of its discovery-set members, and each validation subject was assigned to the nearest exemplar.

To determine whether sex dominance observed in the patient subtypes reflected psychosis-related heterogeneity rather than sex differences already present in the healthy brain, the 640 CNs were additionally assigned to the subtype representations derived from the discovery patient set, using the same assignment procedure as for the validation subjects.

### Bayesian sex test for clustering

After clustering, we assessed whether each resulting cluster was male-dominant, female-dominant, or showed no sex dominance. The natural test is to compare a resulting cluster’s male-to-female ratio against that of the input background (supplied to the algorithm) through a binomial test. This approach, however, is sensitive to the demographic imbalance already present in the input population, as cluster-level sex dominance is evaluated relative to the background distribution. Such a test also does not transfer across datasets, because anchoring a new dataset to the existing cohort with different sex compositions will yield p-values on incommensurable scales.

To address this, we developed a Bayesian test that models the sex selectivity of the clustering mechanism directly, rather than deviation from the input background. We model the clustering mechanism as an “includer”: for each resulting cluster, a single includer admits each male and each female into the cluster with its own inclusion probability *(pmale* and Pfemale). Because these two probabilities are defined relative to their own sex pools (either male or female), the input population’s imbalance does not bias them. Testing for a sex difference is then conducted through a Bayesian model comparison between an includer with a single shared probability (no difference: *pmale* = p_female_) and one with two separate probabilities (difference: *pmale* ≠ p_female_). Because the likelihood integrates against its conjugate prior in closed form, all quantities are computed analytically, without sampling (e.g., MCMC or Gibbs sampling) or approximation (e.g., Laplace or variational methods).

For the discovery set, we test the sex dominance of a given cluster through this two-model comparison: a single-probability includer (no sex dominance) versus a two-probability includer (sex dominance, either male- or female-dominant). When the model posterior, represented as P(dom | D) in Tables 1 and 2, favors the two-probability model (P(dom | D) > 0.5), the cluster is considered as exhibiting sex dominance. To infer the direction of this dominance, we compute the expected male fraction of the cluster under a balanced (50:50) population, represented as P(M) in Tables 1 and 2 (see Supplementary Material, *summary statistics for algorithm-induced sex differences*); P(M) > 0.5 indicates a male-dominant cluster, and P(M) < 0.5 a female-dominant one. For each inclusion probability, we use Jeffreys prior to reflect a noninformative prior regarding sex dominance (see [42]).

**Table 1.** Sex composition and sex dominance of the subtypes recovered by each clustering algorithm. For each algorithm (C-ICA, HYDRA, CHIMERA, K-means, and CSU-RSC) and each subtype, the number of male (M) and female (F) subjects assigned to the cluster is shown for the discovery (N = 992) and validation (N = 247) sets. P(M) is the expected male ratio of the cluster under a perfectly balanced (50:50) population. P(dom | D) is the posterior probability that the cluster is sex-dominant given the data, computed with the Bayesian sex-difference test; for the validation set, discovery posteriors are carried forward as priors, so P(dom | D) quantifies whether the dominance observed in discovery replicates (see Methods). In the discovery set, clusters with strong evidence of sex dominance (P(dom | D) > 0.9) are shown in bold; in the validation set, a cluster is shown in bold only if it was sex-dominant in discovery and its dominance persists (P(dom | D) > 0.5). The direction of dominance is indicated by P(M) (>0.5, male-dominant; < 0.5, female-dominant). Subtype numbering is arbitrary; for readability, male-dominant clusters are placed in the left columns and female-dominant clusters in the right. For CSU-RSC, the target supervised label was set to male for subtypes 1–2 and female for subtypes 3–4, softly biasing each toward the corresponding sex.

|  |  | SUBTYPE 1 | SUBTYPE 2 | SUBTYPE 3 | SUBTYPE 4 |
| --- | --- | --- | --- | --- | --- |
| C-ICA | Discovery | **M = 141, F = 89**<br>**P(M) = 0.61**<br>**P(dom\|D) = 1.00** | M = 135, F = 111<br>P(M) = 0.51<br>P(dom\|D) = 0.17 | M = 124, F = 128<br>P(M) = 0.50<br>P(dom\|D) = 0.05 | **M = 98, F = 166**<br>**P(M) = 0.37**<br>**P(dom\|D) = 1.00** |
|  | Validation | **M = 36, F = 24**<br>**P(M) = 0.56**<br>**P(dom\|D) = 0.61** | M = 40, F = 31<br>P(M) = 0.52<br>P(dom\|D) = 0.51 | M = 32, F = 31<br>P(M) = 0.49<br>P(dom\|D) = 0.48 | **M = 23, F = 30**<br>**P(M) = 0.41**<br>**P(dom\|D) = 0.71** |
| HYDRA | Discovery | M = 102, F = 75<br>P(M) = 0.53<br>P(dom\|D) = 0.35 | M = 90, F = 101<br>P(M) = 0.50<br>P(dom\|D) = 0.07 | M = 187, F = 185<br>P(M) = 0.50<br>P(dom\|D) = 0.05 | M = 119, F = 133<br>P(M) = 0.50<br>P(dom\|D) = 0.08 |
|  | Validation | M = 18, F = 15<br>P(M) = 0.52<br>P(dom\|D) = 0.35 | M = 29, F = 35<br>P(M) = 0.48<br>P(dom\|D) = 0.52 | M = 48, F = 49<br>P(M) = 0.50<br>P(dom\|D) = 0.48 | M = 36, F = 17<br>P(M) = 0.52<br>P(dom\|D) = 0.41 |
| CHIMERA | Discovery | M = 109, F = 105<br>P(M) = 0.50<br>P(dom\|D) = 0.05 | M = 106, F = 94<br>P(M) = 0.50<br>P(dom\|D) = 0.07 | M = 170, F = 180<br>P(M) = 0.50<br>P(dom\|D) = 0.06 | M = 113, F = 115<br>P(M) = 0.50<br>P(dom\|D) = 0.05 |
|  | Validation | M = 33, F = 32<br>P(M) = 0.50<br>P(dom\|D) = 0.43 | M = 22, F = 24<br>P(M) = 0.50<br>P(dom\|D) = 0.39 | M = 40, F = 38<br>P(M) = 0.49<br>P(dom\|D) = 0.47 | M = 36, F = 22<br>P(M) = 0.51<br>P(dom\|D) = 0.49 |
| K-MEANS | Discovery | M = 69, F = 71<br>P(M) = 0.50<br>P(dom\|D) = 0.05 | M = 99, F = 96<br>P(M) = 0.50<br>P(dom\|D) = 0.05 | M = 131, F = 120<br>P(M) = 0.50<br>P(dom\|D) = 0.06 | M = 199, F = 207<br>P(M) = 0.50<br>P(dom\|D) = 0.06 |
|  | Validation | M = 21, F = 16<br>P(M) = 0.50<br>P(dom\|D) = 0.45 | M = 22, F = 26<br>P(M) = 0.50<br>P(dom\|D) = 0.47 | M = 39, F = 24<br>P(M) = 0.52<br>P(dom\|D) = 0.63 | M = 49, F = 50<br>P(M) = 0.49<br>P(dom\|D) = 0.53 |
| CSU-RSC | Discovery | **M = 158, F = 96**<br>**P(M) = 0.62**<br>**P(dom\|D) = 1.00** | M = 144, F = 110<br>P(M) = 0.53<br>P(dom\|D) = 0.48 | M = 104, F = 130<br>P(M) = 0.48<br>P(dom\|D) = 0.28 | **M = 92, F = 158**<br>**P(M) = 0.37**<br>**P(dom\|D) = 1.00** |
|  | Validation | **M = 42, F = 24**<br>**P(M) = 0.60**<br>**P(dom\|D) = 0.87** | M = 45, F = 31<br>P(M) = 0.54<br>P(dom\|D) = 0.66 | M = 24, F = 29<br>P(M) = 0.46<br>P(dom\|D) = 0.65 | **M = 20, F = 32**<br>**P(M) = 0.37**<br>**P(dom\|D) = 0.94** |

**Table 2.** Sex dominance of the patient-derived subtypes when CNs are projected onto them. The patient rows reproduce the discovery-set clustering from Table 1, and the CN rows show the resulting cluster assignments when CNs (N = 640) are assigned to the patient-derived subspaces by minimum reconstruction error (see Methods). The entries (M, F, P(M), P(dom | D)) and the Bayesian sex-difference test are defined as in Table 1. Male- and female-dominant clusters are shown in bold (P(dom | D) > 0.9 for both patients and controls); the direction of dominance is given by P(M) as in Table 1.

|  |  | SUBTYPE 1 | SUBTYPE 2 | SUBTYPE 3 | SUBTYPE 4 |
| --- | --- | --- | --- | --- | --- |
| CSU-RSC | Patient | **M = 158, F = 96**<br>**P(M) = 0.62**<br>**P(dom\|D) = 1.00** | M = 144, F = 110<br>P(M) = 0.53<br>P(dom\|D) = 0.48 | M = 104, F = 130<br>P(M) = 0.48<br>P(dom\|D) = 0.28 | **M = 92, F = 158**<br>**P(M) = 0.37**<br>**P(dom\|D) = 1.00** |
|  | CN | **M = 98, F = 98**<br>**P(M) = 0.60**<br>**P(dom\|D) = 0.95** | M = 59, F = 85<br>P(M) = 0.51<br>P(dom\|D) = 0.25 | **M = 58, F = 114**<br>**P(M) = 0.44**<br>**P(dom\|D) = 0.92** | M = 46, F = 82<br>P(M) = 0.49<br>P(dom\|D) = 0.10 |

For the validation set, we perform a comparison across four includer models: a single probability with (1) a non-informative Jeffreys prior or (2) the discovery-set posterior as prior, and two separate probabilities with (3) a non-informative Jeffreys prior or (4) the discovery-set posteriors as priors. Incorporating the discovery-set posteriors addresses the discovery-dependent question inherent to validation—whether the sex dominance observed in discovery persists in the validation set—while the uninformative models (1, 3) accommodate the case in which the validation set departs substantially from discovery. P(dom | D) is then obtained by Bayesian model averaging over models (2) and (4), and P(M) by averaging over all four models [43, 44]. Full derivations, priors, and closed-form expressions are provided in the Supplementary Material.

### Simulation with SimTB

To validate CSU-RSC against ground truth, we generated resting-state fMRI data using SimTB [45], a MATLAB-based toolbox that simulates fMRI datasets under a model of spatiotemporal separability. The fMRI data are modeled as a linear combination of predefined spatial maps and their associated time courses. We simulated fMRI data comprising 10 spatially ICs (see Figure 2a): three default mode networks (DMNs; medial frontal, precuneus, and angular gyrus; SimTB source ID = 6, 7, 8), two dorsal attention networks (DANs; lateral frontal and inferior parietal sulcus; SimTB source ID = 18, 24), one auditory network (SimTB source ID = 27), two white-matter (SimTB source ID = 16, 17), and two cerebrospinal-fluid components (SimTB source ID = 14, 15). Four subgroups were defined by the spatial spread (network size) of the DMNs and DANs, with all four combinations of large/small DMN and large/small DAN represented. Large networks were 30% larger than their small counterparts.

**Figure 2.**
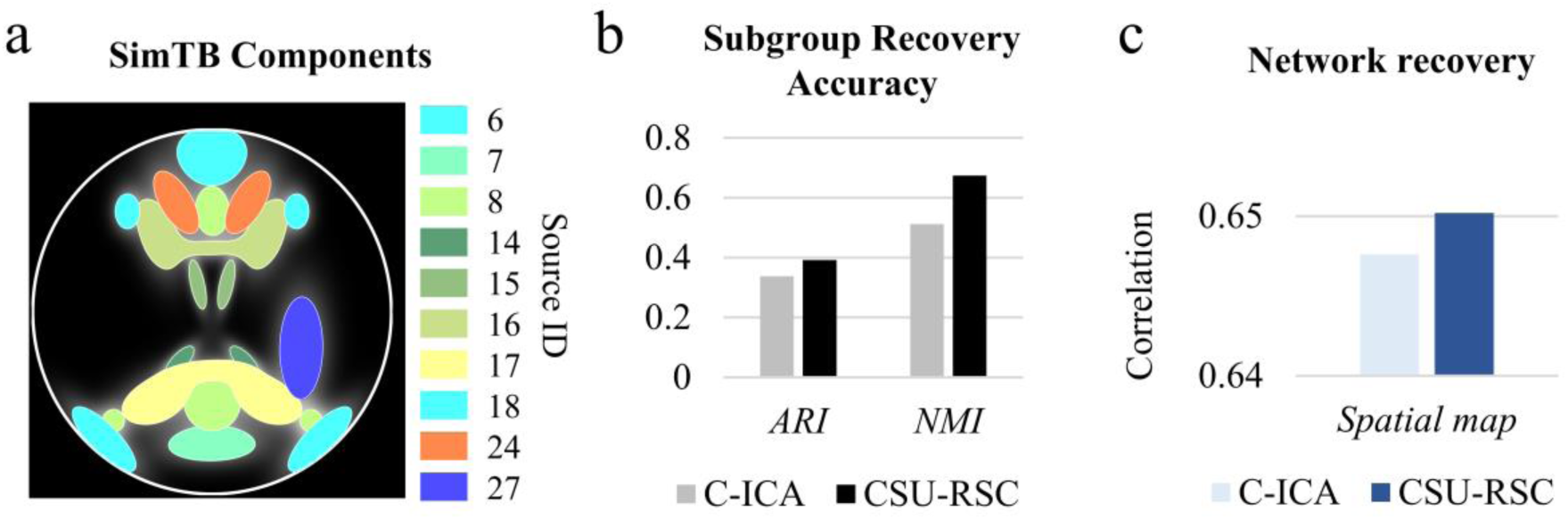
Simulation results. **(a)** The 10 network components (i.e., sources) in SimTB used in fMRI data simulation. **(b)** Accuracy of subgroup recovery, quantified by the adjusted Rand index (ARI) and normalized mutual information (NMI) between the estimated and ground-truth four-subgroup partitions. **(c)** Accuracy of source (network) recovery, quantified by the correlation between the estimated and ground-truth spatial maps. Results are shown for the fully unsupervised C-ICA (light) and the proposed CSU-RSC (dark) in both panels.

Two subgroups were designated male-dominant and two female-dominant, structured so that male-dominant subgroups had large DMNs while female-dominant subgroups had small DMNs, and the two subgroups within each sex differed in DAN size (one large, one small). Each subject was then assigned a sex label such that the dominant sex made up 75% of its subgroup, so that sex was associated with, but not deterministic of, subgroup membership, which mirrors the sex-dominant (rather than sex-exclusive) heterogeneity.

To introduce individual variability, each subject’s network sizes were jittered by a multiplicative factor of 1 + 0.03 • e (where *ε* ∼ *N*(0,1)), i.e., a 3% standard-deviation perturbation around the subgroup mean value. We simulated 50 subjects per subgroup, each with 148 × 148 voxels and 150 time points at a TR of 2s. To mimic the resting state, component time courses were driven solely by unique events, without any task-modulated events.

Recovery of network spatial maps was assessed against the SimTB ground truth. Estimated clusters were first matched to true subgroups by Hungarian algorithm. Within each matched pair, subject-specific spatial maps were reconstructed by spatiotemporal regression [15, 16, 46]. These were compared with each subject’s own ground-truth maps by Hungarian assignment on absolute Pearson correlations.

### Psychosis dataset and preprocessing

We applied the algorithm to real resting-state fMRI data to investigate sex-related heterogeneity in psychosis. The current study included 1,879 participants, comprising 1,239 individuals with a psychotic disorder and 640 CNs. Diagnoses were established using the Structured Clinical Interview for DSM disorders and consensus diagnostic procedures. CNs had no history of psychotic syndromes or recurrent mood syndromes.

All participants underwent a single 5-min resting-state fMRI run on a 3T MRI scanner. During acquisition, they were instructed to remain still, keep their eyes open, and fixate on a crosshair displayed on a monitor. Head motion was further restricted using a custom-built head-coil cushion. Participants’ alertness was assessed immediately after scanning, and the scan was repeated when necessary. fMRI data was preprocessed using the FMRIB Software Library (FSL, version 6.0) and Statistical Parametric Mapping (SPM12) implemented in MATLAB. Head motion was corrected using FSL’s MCFLIRT function, and susceptibility-induced distortions were corrected using FSL’s TOPUP tool. Slice-timing correction was subsequently performed in SPM12. Functional images were spatially normalized to Montreal Neurological Institute (MNI) space using an echo-planar imaging template (i.e., *EPInorm* approach [47]), and resampled to 3 mm isotropic voxels using SPM12. Then, the images were spatially smoothed with a 6 mm FWHM Gaussian kernel.

### Clinical characterization

Clinical characteristics were assessed using four summary measures. Cognitive impairment was evaluated using the Brief Assessment of Cognition in Schizophrenia (BACS) composite score [48], which summarizes performance across verbal memory, digit sequencing, token motor, verbal fluency, symbol coding, and Tower of London tasks. Position along the schizophrenia–bipolar spectrum was characterized using the Schizo-Bipolar Scale (SBS) [49].

Overall psychosis symptom severity was assessed using the Positive and Negative Syndrome Scale (PANSS) total score [50], and social functioning was evaluated using the Birchwood Social Functioning Scale (SFS) [51]. Discovery and validation subjects were combined for these analyses. Prior to group comparisons, age, scanning site, and antipsychotic dose expressed as chlorpromazine (CPZ) equivalents [52] were regressed out of each clinical measure. Subjects with missing data for the respective clinical measure or any covariate were excluded from that analysis. Differences across the four subtypes were assessed using one-way ANOVA, followed by Tukey–Kramer post-hoc comparisons when appropriate.

### Network differences between subtypes

To characterize neurobiological differences among the identified PINs, we compared their subtype-specific ICNs based on spatial similarity. For each pair of subtypes, pairwise spatial Pearson correlations were calculated between their ICNs, and the components were matched using the Hungarian algorithm to maximize the total absolute correlation. Overall similarity between two subtype subspaces was summarized as the mean absolute correlation across the matched ICN pairs. We then examined the matched ICNs of the subtypes individually to identify the networks showing the greatest spatial divergence, after excluding components reflecting motion-related artifacts.

## Results

### Simulation validation

To validate CSU-RSC, we applied it and the fully unsupervised C-ICA to fMRI data simulated with SimTB (see Methods). The simulated data contained four subgroups (N = 50 each; two male-dominant and two female-dominant) defined by network size, and each subject was given a sex label that matched its subgroup’s sex dominance ratio of 75%. We then compared how well CSU-RSC and C-ICA recovered the ground truth subgroup partition.

Both methods performed well above chance (ARI, NMI > 0), but CSU-RSC recovered the partition more accurately than C-ICA (ARI 0.392 vs. 0.338; NMI 0.674 vs. 0.512; Figure 2b). CSU-RSC also better recovered the ground-truth sources used to generate the data, in terms of the spatial maps (correlation 0.650 vs. 0.648; Figure 2c). This gain shows that softly guiding clustering toward a label can improve recovery of the true structure even when the label is only partially informative about the underlying heterogeneity (here, 75% concordant with subgroup membership), supporting the rationale behind CSU-RSC.

### Psychosis spectrum subtyping

We next applied CSU-RSC to real resting-state fMRI data of psychosis individuals, which comprises patients with SZ, SAD, and BP (see Methods). For comparison, we ran five clustering algorithms—CSU-RSC, C-ICA, HYDRA, CHIMERA, and K-means—using a discovery–validation design, in which subtypes were identified in the discovery set and validation subjects were assigned to the resulting discovery-derived subtypes (see Methods). Notably, C-ICA, which used no supervised label, identified one male-dominant and one female-dominant cluster in the discovery set (posterior = 100% each, Bayesian sex test), and this pattern persisted in the validation set (posterior > 50% each; Table 1). In contrast, HYDRA, CHIMERA, and K-means clusters showed no such sex dominance (posterior < 50%). This suggests that sex-associated differences are encoded more explicitly in the brain activity subspaces (i.e., ICN subspaces), compared to the centroid- or boundary-based FNC representations.

Importantly, that this sex structure arises spontaneously, without any access to the labels, indicates that sex is an intrinsic axis of heterogeneity in brain activity. This is precisely the condition under which sex-aware subtyping is warranted: when the signal is present in the data, a method that explicitly incorporates the label can recover it more robustly. Consistent with this, CSU-RSC likewise recovered one male-dominant and one female-dominant cluster (posterior = 100% each in discovery). Cluster membership of sex-dominant clusters showed substantial overlap between C-ICA and CSU-RSC: 94.8% of individuals in the C-ICA male-dominant cluster and 85.8% of those in the CSU-RSC male-dominant cluster are matched; the corresponding overlap was 89.8% and 94.8%, respectively, for the female-dominant clusters. Nonetheless, its validation posteriors were higher than those of the fully unsupervised C-ICA (male-dominant, 87% vs. 61%; female-dominant, 94% vs. 71%), and the expected male fraction (P(M), which quantifies the degree of sex dominance) was more consistent between the discovery and validation sets than for C-ICA (see Table 1), indicating that incorporating the sex label yielded more robust subtypes.

Table 1 reports the full subtyping and sex-test results for all algorithms.

### Specificity of sex dominance to patients

We next examined whether the sex dominance observed in the patient-derived subtypes under CSU-RSC was also present in CNs by assigning CNs to the discovery-derived subtypes (see Methods). The sex-dominant PIN structure did not carry over intact to the healthy population. The female-dominant subspace identified in patients did not reproduce in CNs (posterior = 10%), whereas the male-dominant subspace remained (posterior = 95%; see Table 2). Interestingly, a subspace that had been non-sex-dominant in patients instead became female-dominant in CNs (posterior = 92%). Together, these shifts suggest that sex-related organization of the patient-derived subspaces may reflect psychosis-related variation rather than simply capturing sex differences present in the healthy brain.

### Clinical characterization of the subtypes

Next, we examined clinical differences across subtypes identified by the CSU-RSC using BACS, SBS, PANSS, and SFS in the combined discovery and validation samples, after adjusting confounding variables (e.g., age, CPZ; see Methods). The clusters differed most strongly in cognition (BACS) (CSU-RSC: F(3,635) = 5.83, p = 6×10⁻⁴; one-way ANOVA; Figure 3). The most cognitively impaired cluster was female-dominant, showing lower values than the other three (CSU-RSC, clusters 1 vs. 4, 2 vs. 4, 3 vs. 4: Δ = 0.43, 0.55, 0.47; p = 0.0134, 0.0010, 0.0079; Tukey–Kramer post-hoc tests).

**Figure 3.**
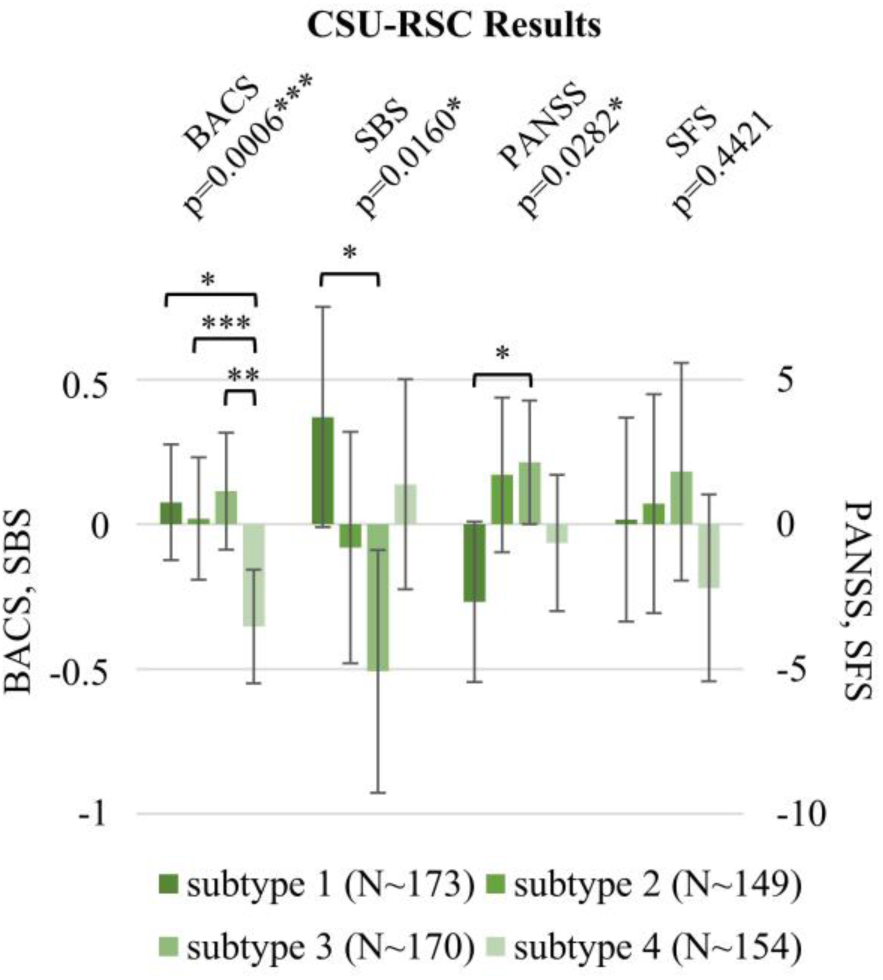
Mean confound-corrected clinical measures for each subtype identified by CSU-RSC, in the combined discovery and validation sets. Age, scanning site, and standardized antipsychotic dose (chlorpromazine equivalents, CPZ) were regressed out of each measure prior to comparison. Clinical scores include BACS composite (cognition), SBS (position along the schizophrenia–bipolar spectrum), PANSS total (symptom severity), and SFS (everyday social functioning). BACS and SBS are plotted against the left axis; PANSS and SFS against the right. Note that higher values indicate better cognition for BACS, a more schizophrenia-like position for SBS, greater symptom severity for PANSS, and better functioning for SFS. Error bars denote 95% confidence intervals (CIs). Values above each score group are one-way ANOVA p-values across the four subtypes; brackets denote pairwise contrasts surviving Tukey–Kramer post-hoc correction (*p < 0.05, **p < 0.01, ***p < 0.001). Legend values give the mean sample size across the four score comparisons for each subtype. Sample sizes vary by measures because individuals missing a confound value or the clinical measure in question were excluded from that comparison. Per-measure sample sizes were, for subtypes 1–4 respectively: (BACS) N = 169, 152, 149, 169; (SBS) N = 176, 157, 179, 153; (PANSS) N = 176, 157, 179, 153; (SFS) N = 170, 129, 173, 140.

SBS also differed significantly across subtypes (CSU-RSC: F = 3.47, p = 0.016; one-way ANOVA), with the male-dominant cluster showing a more schizophrenia-like profile than non-dominant cluster 3 (Δ = 0.88, p = 0.010; Tukey–Kramer). PANSS scores likewise differed significantly across subtypes (F = 3.05, p = 0.028, one-way ANOVA), driven by greater symptom severity in the male-dominant cluster relative to cluster 3 (Δ = 4.82, p = 0.042, Tukey– Kramer). Across the four clinical measures, the significant pairwise differences consistently involved a sex-dominant cluster, suggesting that sex-associated neurobiological heterogeneity captured by CSU-RSC is also clinically relevant.

### Network differences between sex-dominant subtypes

Lastly, we compared the subtype-specific ICNs of the male- and female-dominant CSU-RSC subtypes (see Methods). Despite being distinguished as separate subtypes, they showed the highest overall ICN similarity among all subtype pairs (mean matched-ICA correlation, r = 0.861; Table 3). Among the matched ICNs, the SN showed the greatest spatial divergence after excluding motion-related components (Figure 4a), with the male-dominant subtype showing a sharper contrast between default-mode and task-positive regions (Figure 4b), consistent with SN disruption reported in relation to psychotic symptoms and cognitive dysfunction [53, 54]. Notably, the SNs of the non-sex-dominant subtypes (subtypes 2 and 3) differed markedly from those of the sex-dominant subtypes (Figure 4a), with weakened coherent activity in frontal regions (Figure 4c).

**Figure 4.**
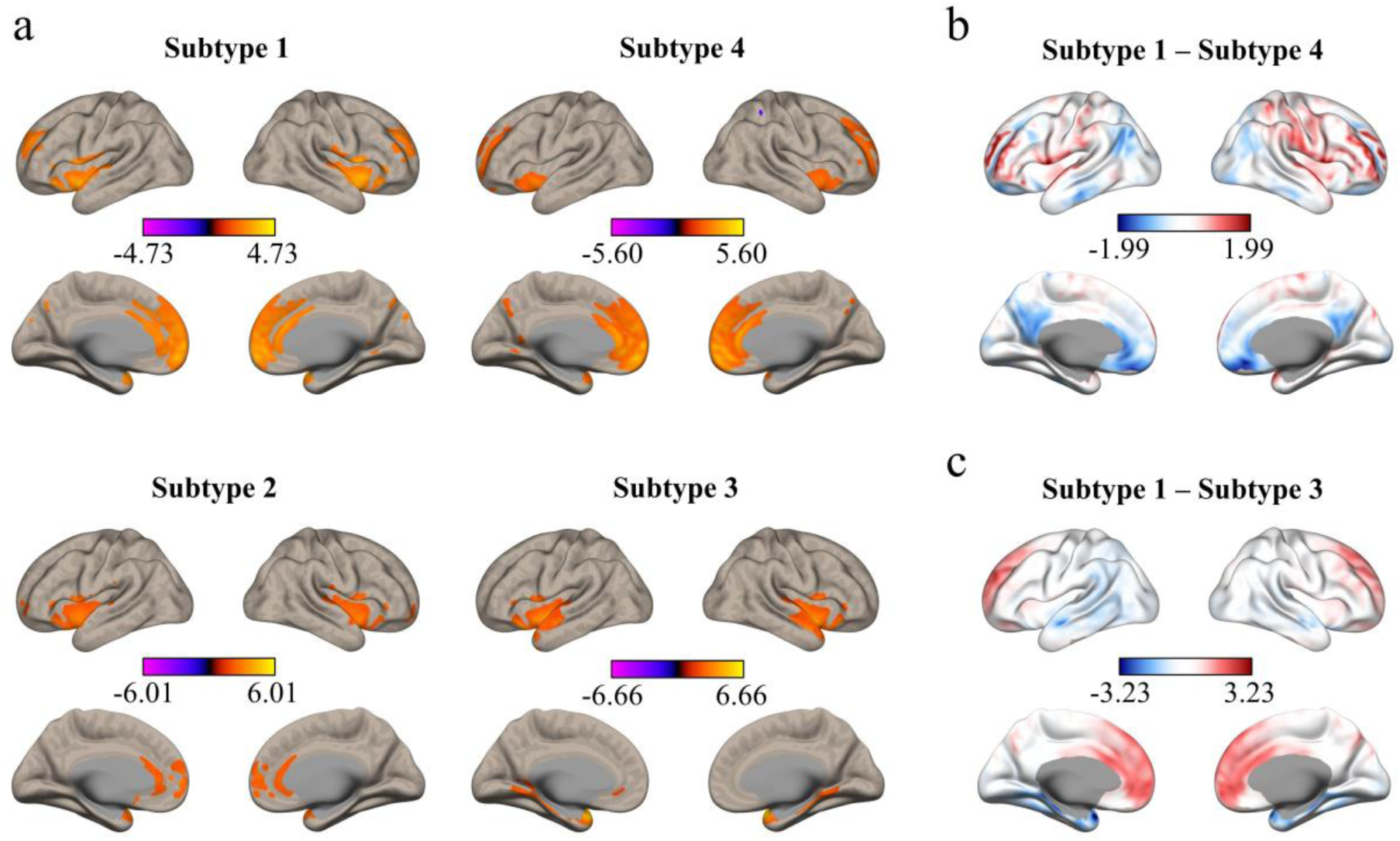
Salience-network (SN) differences across CSU-RSC PINs. **(a)** Salience-network spatial maps for the four CSU-RSC subtypes (subtypes 1, male-dominant; subtypes 4, female-dominant; subtype 2 and 3, non-sex-dominant). The values are z-scored across voxels, thresholded by |z| > 2. Although the overall spatial patterns are broadly similar across subtypes, the sex-dominant subtypes (1 and 4) most closely resemble each other, whereas the non-sex-dominant subtypes (2 and 3) show weaker coherent engagement of frontal regions. **(b)** Voxel-wise difference between the male-dominant (subtype 1) and female-dominant (subtype 4) salience networks (subtype 1 - subtype 4); red indicates higher z-score in subtype 1, blue regions higher in subtype 4. **(c)** Voxel-wise difference between subtype 1 and the non-sex-dominant subtype 3 (subtype 1 - subtype 3), shown for comparison; red indicates higher z-score in sex-dominant subtype (i.e., subtype 1), blue regions higher in the non-sex-dominant subtype (i.e., subtype 3).

**Table 3.** Mean absolute Pearson correlation between subtype-specific ICNs identified by CSU-RSC. For each subtype pair, components were matched one-to-one using the Hungarian algorithm to maximize total absolute correlation, and the values shown are averaged across matched pairs.

|  | SUBTYPE 1 | SUBTYPE 2 | SUBTYPE 3 | SUBTYPE 4 |
| --- | --- | --- | --- | --- |
| SUBTYPE 1 |  | 0.6438 | 0.6264 | 0.8614 |
| SUBTYPE 2 |  |  | 0.5687 | 0.6263 |
| SUBTYPE 3 |  |  |  | 0.6834 |
| SUBTYPE 4 |  |  |  |  |

## Discussion

In this study, we introduced CSU-RSC, a framework that incorporates a supervised label into subspace-based clustering with applications to neuroimaging data. By softly guiding the partitioning toward label-relevant structure (here, sex), rather than ignoring the label or enforcing it as a hard constraint, CSU-RSC aims to more precisely recover label-related heterogeneity in neurobiology. On simulated data with known ground truth, CSU-RSC recovered the underlying subgroups more accurately than its fully unsupervised counterpart— even when the label was only partially aligned with the true partition. Applied to psychosis, it identified sex-dominant neurosubtypes (i.e., sex-dominant PINs) validated by differential clinical characteristics across cognitive and symptom dimensions.

A key observation is that sex-dominant PINs emerged even with the fully unsupervised method (i.e., C-ICA), with sex labels completely unknown, which provides independent evidence for substantial sex-related heterogeneity in intrinsic brain organization. Notably, this structure emerged only with the subspace-based methods, not with the centroid- or boundary-based approaches (i.e., HYDRA, CHIMERA, and K-means) using FNC features, suggesting that sex-associated variation may be more readily captured in brain activity subspaces. As our simulation indicated that this is precisely the setting in which label-aware subtyping can be beneficial. Consistent with this speculation, CSU-RSC yielded more robust PINs, with sex dominance more consistently preserved in the validation set (see Results).

These sex-dominant PINs did not merely reflect demographic segregation. The subtypes differed along distinct clinical axes: cognitive performance, position on the schizophrenia– bipolar spectrum, and symptom severity. In each case, the clinical heterogeneity was more carried by the sex-dominant ones. The most cognitively impaired subtype (lowest BACS) was female-dominant; the most schizophrenia-like position on the spectrum (high SBS) was expressed in a male-dominant subtype; and the male-dominant subtype showed the greatest symptom severity (highest PANSS). The greater symptom severity and more schizophrenia-like spectrum position of the male-dominant subtype are consistent with previous reports that male patients tend toward earlier onset, more severe symptoms, and a more typical schizophrenia profile [32, 33, 55]. In contrast, the cognitive finding runs counter to the frequently reported greater cognitive impairment in male patients, although evidence for sex differences in cognitive impairment remains mixed in psychosis [56, 57].

At the network level, while the male- and female-dominant CSU-RSC PINs were distinguished by detectable differences in their subspaces, they nevertheless showed the highest mean matched-ICN correlation among all subtype pairs. The SN showed the greatest divergence between the two, consistent with its established involvement in psychosis-related symptoms and cognitive dysfunction [53, 54, 58], although its relevance to the observed subtype differences requires further investigation.

The CN assignment results offer an important insight, helping distinguish sex dominance that is disorder-specific from sex dominance that reflects general sex differences. The two sex-dominant PINs diverged in this respect. The male-dominant subspace remained male-dominant in CNs, suggesting it partly reflects general sex differences in the healthy brain’s functional organization (e.g., [59, 60]). In contrast, the female-dominant subspace did not remain female-dominant in CNs, suggesting that its sex dominance may be more closely related to psychosis. A female dominance instead emerged in the subspace of subtype 3, which had been non-dominant and comparatively bipolar-leaning in patients. This might reflect a female-linked affective vulnerability [61–63], though the interpretation warrants caution, as it could equally arise from subjects displaced from the female-dominant subspace under the current hard assignment (i.e., assignment to a single cluster that minimizes reconstruction error). Nonetheless, taken together, these results suggest that subspace-based clustering may help distinguish disorder-related sex-associated heterogeneity from normative sex differences in healthy brain organization (see [64, 65]).

Lastly, we emphasize that CSU-RSC is a general framework for more accurate neurosubtyping rather than a single algorithm. Its supervised label is interchangeable—here we use sex, but equally diagnosis, age, genetic risk, or any categorical axis expected to structure the heterogeneity can be used—and the framework is agnostic to the clustering method, extending beyond the subspace-based decomposition we used to general clustering approaches. The degree of supervision is self-tunable. In the present implementation, we controlled it through the dropout in refinement step (Step 4; see Figure 1), but the framework readily admits additional controls—e.g., biasing the reassignment step (Step 3) toward the supervised label— which together could span a continuum from fully unsupervised to fully label-stratified clustering. A comprehensive investigation of how these parameter choices trade off against subtyping accuracy is beyond the scope of a single cross-sectional study and represents an important direction for future research. By establishing both the framework and the methodological tools for its further development, CSU-RSC offers a foundation for label-aware neurosubtyping.

## Code availability

All analyses in this study were conducted using MATLAB R2025a. The scripts and code supporting the study’s findings are available on GitHub and will be publicly accessible upon the paper’s publication.

## Acknowledgement

This work was supported by the National Institutes of Health grant numbers R01MH136665 (to A.I. and J.C.).

