## Supplementary Material for "Concurrent Supervised-Unsupervised Representative Subspace Clustering (CSU-RSC)"

**Bayesian Sex Difference Test**

*Setup & Notation*

Consider a clustering algorithm that partitions a background input population into clusters. We examine one cluster at a time. Let the background population contain $M$ males and $F$ females, of which the algorithm assigns $m$ males and $f$ females to the cluster. We model the clustering mechanism as an *includer* that admits each male into the cluster independently with probability $p_m$ and each female independently with probability $p_f$. The probabilities of observing $m$ males and $f$ females in the cluster then follow binomial distributions:

$$p(m) = \binom{M}{m} p_m^m (1 - p_m)^{M-m}$$

$$p(f) = \binom{F}{f} p_f^f (1 - p_f)^{F-f}$$

Note that because $p_m$ and $p_f$ are each defined for each sex, any difference between them reflects the sex selectivity of the algorithm itself, independent of the sex-ratio imbalance in the input population. See Supplementary Figure 1.

Testing whether a given cluster is male- or female-dominant therefore reduces to asking whether the clustering algorithm treats the two sexes differently, i.e., to comparing the following two models:

- $\mathcal{M}_1$**(no sex dominance):** $p_m = p_f = p$, with $p \sim \text{Beta}(0.5,0.5)$.
- $\mathcal{M}_2$**(sex dominance):** $p_m$, $p_f$ independent, with $p_m$, $p_f \sim \text{Beta}(0.5,0.5)$.

Here, $\text{Beta}(0.5,0.5)$ is the Jeffreys prior, which less influence the posterior compared to the uniform prior $\text{U}([0,1])$ [1].

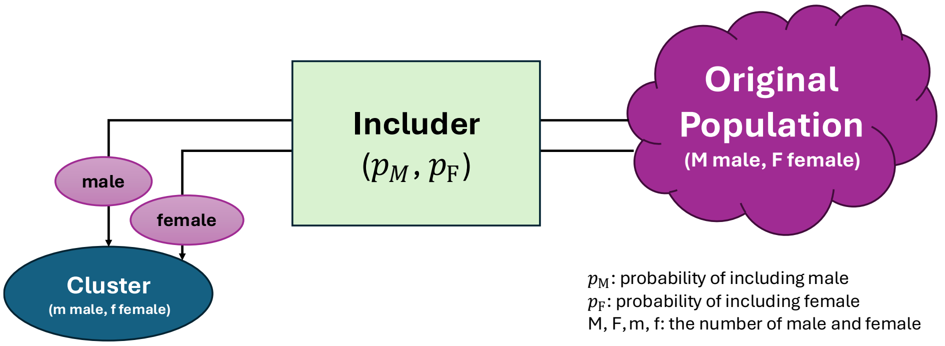


**Supplementary Figure 1.** Includer model.

*Bayesian model comparison*

Since testing for sex dominance reduces to a model comparison, its statistical significance can be assessed within a Bayesian framework. Let $D = (m, f)$ concisely denote the observed clustering result. The *likelihood* of observing $D$ in each model is given by the definition of each model (i.e., $\mathcal{M}_1$ and $\mathcal{M}_2$), which is a multiplication of the probabilities of observing $m$ males and $f$ females in the cluster ($p(m)$ and $p(f)$ in above).

Under $\mathcal{M}_1$ (no sex dominance), the likelihood is

$$P(D|\mathcal{M}_1, p) = \binom{M}{m} p^m (1-p)^{M-m} \binom{F}{f} p^f (1-p)^{F-f}.$$

Under $\mathcal{M}_2$ (sex dominance), the likelihood is

$$P(D|M_2, p_m, p_f) = \binom{M}{m} p_m^m (1-p_m)^{M-m} \binom{F}{f} p_f^f (1-p_f)^{F-f}.$$

For each model, the *marginal likelihood* (or model evidence) is obtained by integrating the likelihood over the prior on its parameters:

$$P(D|\mathcal{M}_1) = \int_0^1 P(D|\mathcal{M}_1, p)\, P(p \mid \mathcal{M}_1) dp$$

$$P(D|\mathcal{M}_2) = \int_0^1 \int_0^1 P(D|M_2, p_m, p_f)\, P(p_m \mid \mathcal{M}_2)\, P(p_f \mid \mathcal{M}_2)\, dp_m dp_f$$

where $P(p|\mathcal{M}_1)$, $P(p_m|\mathcal{M}_2)$, $P(p_f|\mathcal{M}_2)$ are prior probabilities of $p$, $p_m$, and $p_f$.

Then, the two models can be compared through the *Bayes factor*, the ratio of their model evidence,

$$BF_{21} = \frac{P(D|\mathcal{M}_2)}{P(D|\mathcal{M}_1)},$$

which quantifies the relative support the data provide for the sex-difference model over the no-difference model.

Under equal prior probabilities on the two models, the model posterior follows directly as $P(\mathcal{M}_2|D) = BF_{21}/(BF_{21} + 1)$.

*Two-step procedure (transferring knowledge from discovery to validation)*

Because the binomial likelihood and a Beta prior (including the uniform and Jeffreys priors) are conjugate distributions, the marginal likelihoods can be expressed in closed form (each integral reduces to a Beta function), and the model comparison here admits exact analytic solutions. Thus, no sampling strategy (e.g., MCMC or Gibbs sampling) or approximation method (e.g., Laplace approximation or variational Bayesian approach) is required. Below, we give the closed-form posterior expressions for the discovery–validation setting described in the main text.

***Step 1: Discovery.***

Let the discovery clustering results $D_{\text{disc}} = (m_d, f_d)$, in the given cluster among background discovery population with ($M_d$ male, $F_d$ female). Under the Jeffreys prior (see above), the marginal likelihoods take the closed forms:

$$P(D_{\text{disc}}|\mathcal{M}_1) = C_d \frac{\text{Beta}(m_d + f_d + 0.5, (M_d + F_d) - (m_d + f_d) + 0.5)}{\text{Beta}(0.5, 0.5)}$$

$$P(D_{\text{disc}}|\mathcal{M}_2) = C_d \frac{\text{Beta}(m_d + 0.5, M_d - m_d + 0.5) \cdot \text{Beta}(f_d + 0.5, F_d - f_d + 0.5)}{\text{Beta}(0.5, 0.5)^2}$$

where $C_d = \binom{M_d}{m_d}\binom{F_d}{f_d}$ and $\mathrm{Beta}(\alpha, \beta)$ is the Beta function. The Bayes factor and the model posterior (denoted as P(dom | D) in the main text) can be calculated directly from these quantities.

The parameter posteriors $P(p|\mathcal{M}_1; D_{\mathrm{disc}})$, $P(p_m|\mathcal{M}_2; D_{\mathrm{disc}})$, and $P(p_f|\mathcal{M}_2; D_{\mathrm{disc}})$, which describe the plausible values of the inclusion probabilities after observing the discovery clustering result, are likewise available in closed form:

$$P(p|\mathcal{M}_1; D_{\mathrm{disc}}) = \mathrm{Beta}(m_d + f_d + 0.5, (M_d + F_d) - (m_d + f_d) + 0.5)$$

$$P(p_m|\mathcal{M}_2; D_{\mathrm{disc}}) = \mathrm{Beta}(m_d + 0.5,\ M_d - m_d + 0.5)$$

$$P(p_f|\mathcal{M}_2; D_{\mathrm{disc}}) = \mathrm{Beta}(f_d + 0.5,\ F_d - f_d + 0.5)$$

***Step 2: Validation (replication).***

As noted in the main text, the question asked of the validation set is inherently conditioned on the discovery set: whether the sex dominance observed in discovery persists in the validation set. Accordingly, model comparison for the validation set was performed over four includer models: a single probability with (1a) a non-informative Jeffreys prior or (1b) the discovery-set posterior as prior, and two separate probabilities with (2a) a non-informative Jeffreys prior or (2b) the discovery-set posteriors as priors. The marginal likelihoods and parameter posteriors for these four models all take close forms.

Let $D_{val} = (m_v, f_v)$ denote the validation assignment result for the given cluster among background validation population with $M_v$ males and $F_v$ females. For brevity, we write the discovery-set posterior parameters (see above) as

$$\alpha_m = m_d + 0.5, \qquad \beta_m = M_d - m_d + 0.5, \qquad \alpha_f = f_d + 0.5, \qquad \beta_f = F_d - f_d + 0.5,$$

$$\alpha_0 = m_d + f_d + 0.5, \qquad \beta_0 = (M_d + F_d) - (m_d + f_d) + 0.5,$$

and $C_v = \binom{M_v}{m_v}\binom{F_v}{f_v}$ is a common binomial coefficient.

Then, the marginal likelihoods and parameter posteriors are given as follows:

**Model 1a** ($\mathcal{M}_{1a}$: no sex dominance, i.e., single inclusion probability; Jeffreys prior).

- Marginal likelihood: $P(D_{val} \mid \mathcal{M}_{1a}) = C_v \, \frac{Beta(m_v+f_v+0.5, M_v+F_v-m_v-f_v+0.5)}{Beta(0.5,0.5)}$

- Posterior: $P(p|\mathcal{M}_{1a}; D_{\text{val}}) = Beta(m_v + f_v + 0.5, M_v + F_v - m_v - f_v + 0.5)$

**Model 1b** ($\mathcal{M}_{1b}$ no sex dominance, i.e., single inclusion probability; discovery-set posterior as prior).

- Marginal likelihood: $P(D_{val} \mid \mathcal{M}_{1b}; D_{\text{disc}}) = C_v \, \frac{Beta(\alpha_0+m_v+f_v, \beta_0+M_v+F_v-m_v-f_v)}{Beta(\alpha_0,\beta_0)}$

- Posterior: $P(p|\mathcal{M}_{1b}; D_{\text{val}}, D_{\text{disc}}) = Beta(\alpha_0 + m_v + f_v, \beta_0 + M_v + F_v - m_v - f_v)$

**Model 2a** ($\mathcal{M}_{2a}$: sex dominance, i.e., two inclusion probabilities; Jeffreys prior).

- Marginal likelihood: $P(D_{val} \mid \mathcal{M}_{2a}) = C_v \, \frac{Beta(m_v+0.5, M_v-m_v+0.5)\; B(f_v+0.5, F_v-f_v+0.5)}{Beta(0.5,0.5)^2}$

- Posteriors: $P(p_m|\mathcal{M}_{2a}; D_{\text{val}}) = Beta(m_v + 0.5, M_v - m_v + 0.5)$ and

$$P(p_f|\mathcal{M}_{2a}; D_{\text{val}}) = Beta(f_v + 0.5, F_v - f_v + 0.5).$$

**Model 2b** ($\mathcal{M}_{2b}$; sex dominance, i.e., two inclusion probabilities; discovery-set posteriors as priors). That is, $p_m \sim Beta(\alpha_m, \beta_m)$ and $p_f \sim Beta(\alpha_f, \beta_f)$.

- Marginal likelihood: $P(D_{val} \mid \mathcal{M}_{2b}) = C_v \, \frac{Beta(\alpha_m+m_v, \beta_m+M_v-m_v)}{Beta(\alpha_m,\beta_m)} \cdot \frac{Beta(\alpha_f+f_v, \beta_f+F_v-f_v)}{Beta(\alpha_f,\beta_f)}$

- Posteriors: $P(p_m|\mathcal{M}_{2b}; D_{\text{val}}, D_{\text{disc}}) = Beta(m_d + m_v + 0.5, (M_d - m_d) + (M_v - m_v) + 0.5)$ and $P(p_f|\mathcal{M}_{2a}; D_{\text{val}}, D_{\text{disc}}) = Beta(f_d + f_v + 0.5, (F_d - f_d) + (F_v - f_v) + 0.5)$.

Assigning equal prior probability to the four models (= 0.25), the posterior probability of model $\mathcal{M}_i$ given the validation data is

$$P(\mathcal{M}_i \mid D_{val}) = \frac{P(D_{val} \mid \mathcal{M}_i)}{\sum_j P\left(D_{val} \mid \mathcal{M}_j\right)}, \qquad i \in \{\mathcal{M}_{1\mathrm{a}}, \mathcal{M}_{1\mathrm{b}}, \mathcal{M}_{2\mathrm{a}}, \mathcal{M}_{2\mathrm{b}}\}.$$

The model posterior of sex dominance (i.e., P(dom | D)) for the validation set is obtained by Bayesian model averaging over the two sex-dominance models, P(dom | D) = $P(\mathcal{M}_{2a} \mid D_{val})$ + $P(\mathcal{M}_{2b} \mid D_{val})$.

*Summary statistics for algorithm-induced sex differences*

Because we obtain the posterior distributions of the model parameters, we can quantify the genuine sex bias induced by the clustering algorithm. Specifically, we define this bias as the expected male fraction within a cluster under a hypothetical sex-balanced population (M = F) in the large-sample limit ($M, F \to \infty$). This quantity can be estimated as $P(\mathrm{M}) = \hat{p}_m/(\hat{p}_m + \hat{p}_f)$, where $\hat{p}_m$ and $\hat{p}_f$ are posterior means of inclusion probabilities. The mean of $Beta(\alpha, \beta)$ is given by $\alpha/(\alpha + \beta)$. This statistic provides an interpretable measure of algorithm-induced sex dominance: i.e., P(M) > 0.5 indicates a male-dominant cluster, and P(M) < 0.5 a female-dominant one. Moreover, this can be directly compared across datasets (e.g., discovery vs. validation).
